# The Tenets of Teneurin: Conserved Mechanisms Regulate Diverse Developmental Processes in the *Drosophila* Nervous System

**DOI:** 10.1101/499657

**Authors:** Alison T. DePew, Michael A. Aimino, Timothy J. Mosca

## Abstract

To successfully integrate a neuron into a circuit, a myriad of developmental events must occur correctly and in the correct order. Neurons must be born and grow out towards a destination, responding to guidance cues to direct their path. Once arrived, each neuron must segregate to the correct sub-region before sorting through a milieu of incorrect partners to identify the correct partner with which they can connect. Finally, the neuron must make a synaptic connection with their correct partner; a connection that needs to be broadly maintained throughout the life of the animal while remaining responsive to modes of plasticity and pruning. Though many intricate molecular mechanisms have been discovered to regulate each step, recent work showed that a single family of proteins, the Teneurins, regulates a host of these developmental steps in *Drosophila* – an example of biological adaptive reuse. Teneurins first influence axon guidance during early development. Once neurons arrive in their target regions, Teneurins enable partner matching and synapse formation in both the central and peripheral nervous systems. Despite these diverse processes and systems, the Teneurins use conserved mechanisms to achieve these goals, as defined by two tenets: 1) transsynaptic interactions with each other and 2) membrane stabilization via an interaction and regulation of the cytoskeleton. These processes are further distinguished by 1) the nature of the transsynaptic interaction – homophilic interactions (between the same Teneurins) to engage partner matching and heterophilic interactions (between different Teneurins) to enable synaptic connectivity and the proper apposition of pre- and postsynaptic sites and 2) the location of cytoskeletal regulation (presynaptic cytoskeletal regulation in the CNS and postsynaptic regulation of the cytoskeleton at the NMJ). Thus, both the roles and the mechanisms governing them are conserved across processes and synapses. In this review, we will highlight the contributions of *Drosophila* synaptic biology to our understanding of the Teneurins and discuss the mechanistic conservation that allows the Teneurins to achieve common neurodevelopmental goals. Finally, we will posit the next steps for understanding how this remarkably versatile family of proteins functions to control multiple distinct events in the creation of a nervous system.

## INTRODUCTION

In the nervous system, each neuron undergoes a simultaneously elegant yet complex development. Disparate molecular, cellular, and morphological events are woven together into a united entity, linking cell birth, neuronal differentiation, cell migration, membrane adhesion, synapse formation, and synaptic refinement. These diverse processes, with their distinctive molecular, developmental, and cell biological requirements, are united by a common, broad goal: forming the functional connections essential for life. When the diversity of neuronal subtype in different brain regions, layers, and even systems (peripheral versus central) is added to this already herculean list, it becomes apparent that achieving proper development is no easy task. Each process has its own distinct molecular and physical requirements and challenges, and these processes need to be seamlessly connected both spatially and temporally. If these events occur in the wrong order or in the wrong place, development can go awry, resulting in intellectual disabilities and neurodevelopmental disorders including autism, schizophrenia, and bipolar disorder (Gilbert and Man, 2017; Grant, 2012; Zerbi et al., 2018). Thus, the underlying processes must be finely tuned to ensure fidelity in neurodevelopment.

How are these disparate tasks accomplished? Based on estimations of neuronal diversity (Lodato and Arlotta, 2015), genome sizes (Adams et al., 2000; Cravchik et al., 2001), and synapse number in the brain (Silbereis et al., 2016), it would be impossible to employ a different approach with distinct molecular cues and mechanistic underpinnings for each event. This would require more distinct adhesion and recognition cues than there are actual genes in the genome. There must be some shared use of molecules and processes. Indeed, this is commonly observed throughout development where different classes of neurons use similar molecules and mechanisms to accomplish the goals of axon guidance (Dickson, 2002), synapse formation (Favuzzi and Rico, 2018), and neuronal migration (Geschwind and Rakic, 2013). We see this concept in our cities frequently, in the form of “adaptive reuse”: a decommissioned water pumping station becomes a gastropub, a turn-of-the-century bank becomes a museum, and even a former firehouse becomes a luxury apartment complex. This process of using an “old” molecule or concept for a purpose other than its original intent enables considerable utility. At the molecular level, we see genes originally intended for cell adhesion adaptively reused to form synapses (Giagtzoglou et al., 2009; Sun and Xie, 2012) and cytoskeletal molecules used for movement repurposed for cell migration (Etienne-Manneville, 2013). Backed by this concept, the list of distinct processes needed for neuronal development becomes more manageable, as does its molecular requirements.

Recent years have seen an explosion of research on a family of large cell surface proteins called the Teneurins (Young and Leamey, 2009) that play diverse roles in organismal development (Tucker and Chiquet-Ehrismann, 2006). In the fruit fly, *Drosophila melanogaster*, the two Teneurin homologues, Ten-m and Ten-a, were originally thought to be involved in body segment patterning (Baumgartner and Chiquet-Ehrismann, 1993; Baumgartner et al., 1994; Levine et al., 1994; Rakovitsky et al., 2007). The last decade, however, has seen the discovery of roles for these cell surface proteins in multiple neurodevelopmental processes including axon guidance, synaptic partner matching, and synapse organization (Hong et al., 2012; Mosca and Luo, 2014; Mosca et al., 2012; Zheng et al., 2011). *Drosophila* has proven an outstanding model system to assess Teneurin function in that its many genetic tools (Venken et al., 2011), accessible synapses at the NMJ (Harris and Littleton, 2015) and in the olfactory system (Mosca and Luo, 2014) and its stereotyped wiring (Couto et al., 2005; Keshishian et al., 1996) enable detailed molecular and mechanistic study at the single-cell level. In such discovery, a theme of adaptive reuse surfaced for the Teneurins: the same genes controlling multiple steps of neurodevelopment via similar mechanisms. We will focus on two of these processes: synaptic partner matching and synaptic organization to describe recent work highlighting roles for the *Drosophila* Teneurins in both these processes as well as their shared mechanistic underpinnings.

## MATERIALS AND METHODS

### Drosophila Genetics

All stocks and crosses were raised on standard cornmeal / dextrose medium at 25°C in a 12/12 light/dark cycle. Canton S. served as the control strain (Woodard et al., 1989). *Df (X) ten-a* was used as a *ten-a* null mutant (Mosca et al., 2012). *Mef2-GAL4* was used to drive expression in all muscles (Lilly et al., 1995). *SG18.1-GAL4* was used to drive expression in all ORNs (Shyamala and Chopra, 1999). We also used the transgenic strains *UAS-Ten-a* (Mosca et al., 2012) and *UAS-ten-m*^*RNAi-V51173*^ (Mosca et al., 2012) for Ten-a expression and *ten-m* RNAi knockdown, respectively.

### Staining, Spaced Stimulation, and Immunocytochemistry

Spaced stimulation was conducted as previously described (Piccioli and Littleton, 2014). Wandering third instar larvae were processed for immunocytochemistry as previously described (Mosca and Schwarz, 2010). The following primary antibodies were used: mouse anti-Ten-m at 1:500 (Levine et al., 1994), rabbit anti-Dlg at 1:40000 (Koh et al., 1999), rabbit anti-Syt I at 1:4000 (Mackler et al., 2002). Alexa488- and Alexa546-conjugated secondary antibodies were used at 1:250 (Jackson ImmunoResearch and Invitrogen). Cy5-conjugated antibodies to HRP were used at 1:100 (Jackson ImmunoResearch).

### Olfactory Behavior Trap

Olfactory behavior experiments were conducted and analyzed as previously described (Mosca et al., 2017).

### Genotypes

Figure 3: *Control* (*+; +; +; +*); *ten-a -/-* (*Df (X) ten-a; +; +; +*); *ten-a -/-* + ORN Ten-a (*Df (X) ten-a; SG18.1-GAL4 / UAS-Ten-a; +; +*). Figure 4A,C (*+; +; +; +*); Figure 4B (*+; Mef2-GAL4 / +; UAS-ten-m*^*RNAi-V51173*^ */ +; +*).

**Figure 3.**
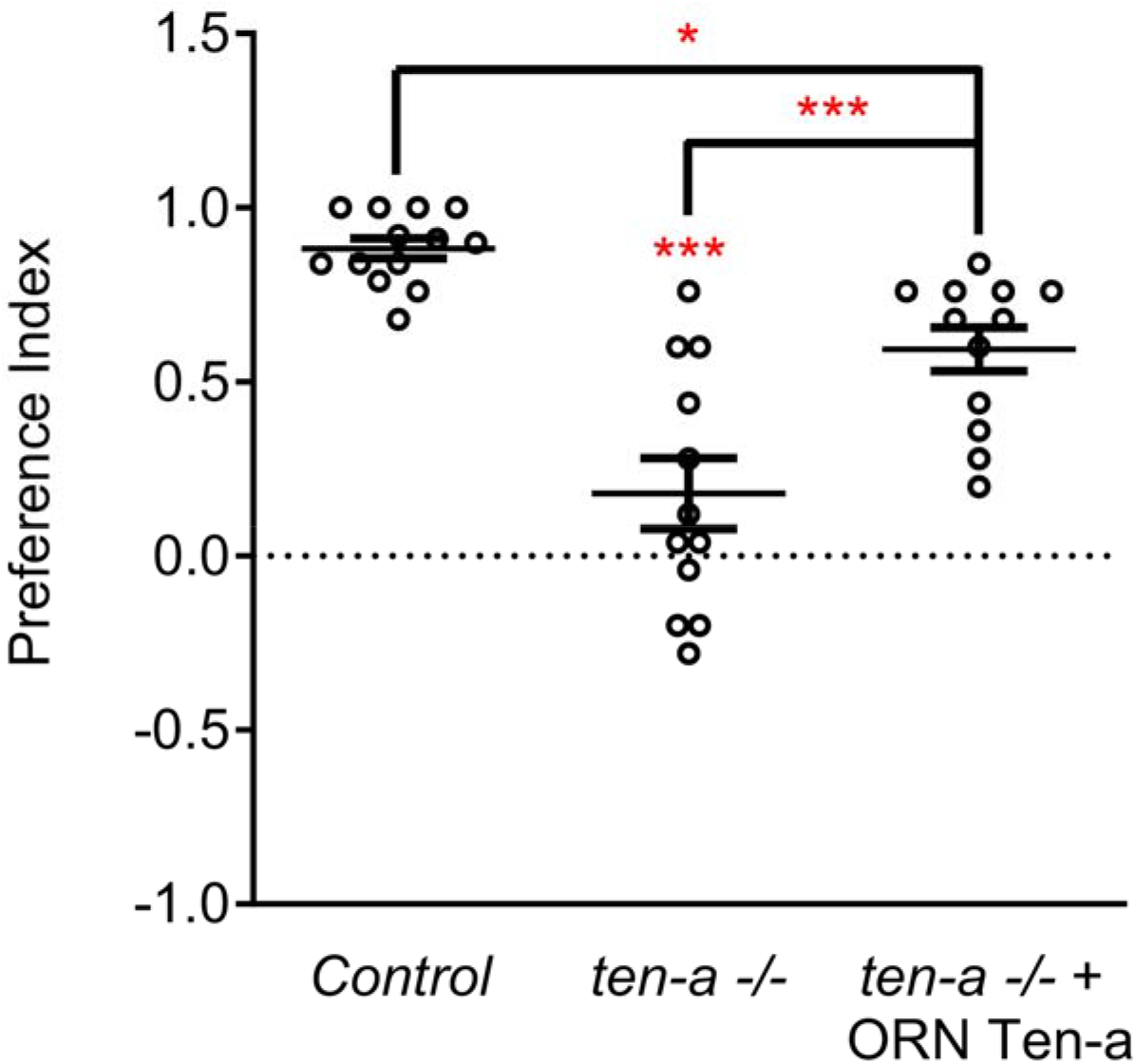
Ten-a is required ORNs for normal odorant attraction. Graph of preference indices for various genotypes in an olfactory trap assay pairing apple cider vinegar (an attractive odorant) and water (a null solution). Control flies exhibit strong attraction to apple cider vinegar while loss of *ten-a* nearly completely abrogates this attraction. This phenotype can be partially rescued by restoring Ten-a expression in ORNs of the *ten-a* mutant, demonstrating that presynaptic Ten-a is required for normal olfactory behavior. N ≥ 12 cohorts of 25 flies each for all experiments. * *p* < 0.05, *** *p* < 0.001.

**Figure 4.**
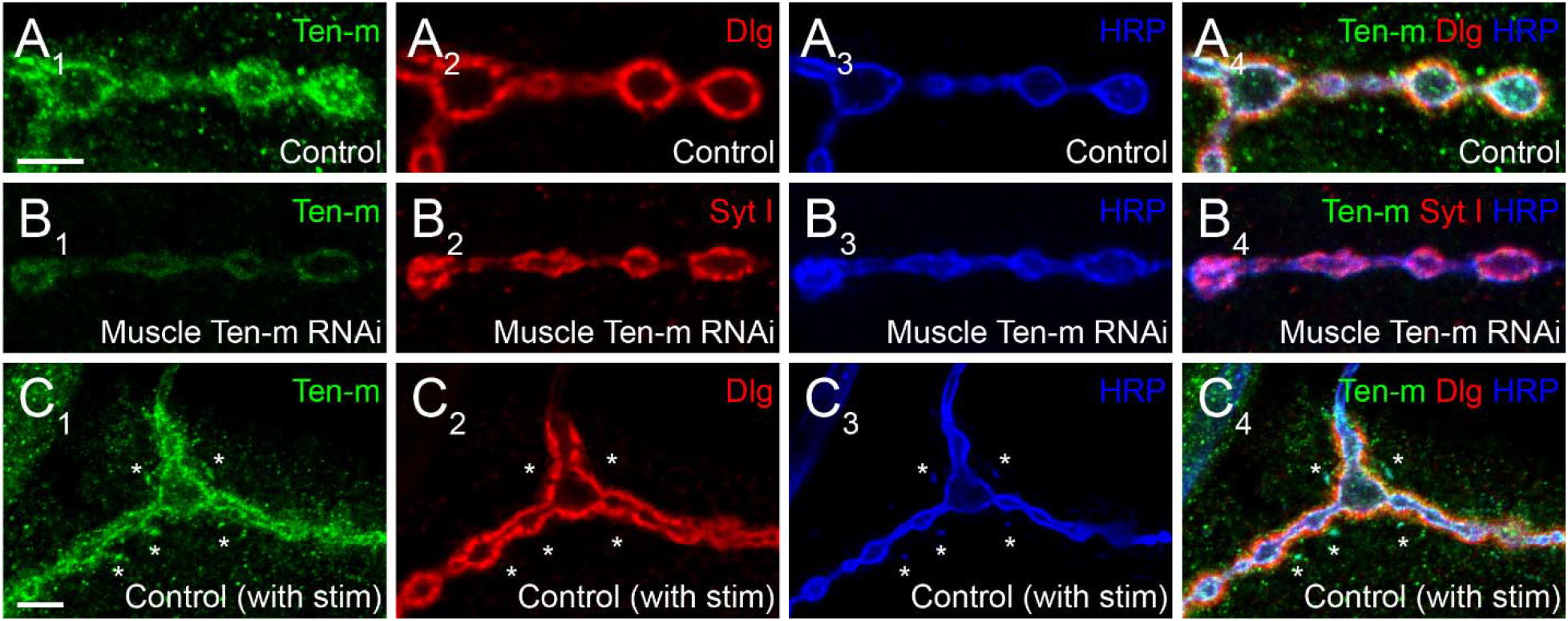
Presynaptic Ten-m localizes to newly formed synaptopods. Single confocal sections of NMJs stained with antibodies to Ten-m (green), Dlg (red), and HRP (blue, in A and C) or Ten-m (green), Syt I (red), and HRP (blue, in B). Control NMJs show predominantly postsynaptic Ten-m localization (A) but a presynaptic component associated with the HRP-positive membrane and synaptic vesicle (via Syt I) population is visible upon genetic removal of the postsynaptic pool of Ten-m (B). Following spaced stimulation, Ten-m (green) is visible within newly formed synaptopods (marked by asterisks) that have not yet been apposed by postsynaptic Dlg (red) staining. Scale = 5 μm.

## PARTNER MATCHING

Before any synapse can be made and organized, the pre- and postsynaptic cells must first identify each other as appropriate partners and begin connecting in a process called partner matching. While there has been extensive research done on many different aspects of synapse formation, the step of partner matching remains poorly understood. In 1963, Roger Sperry proposed that such a process may occur by ‘individual identification tags, presumably cytochemical in nature’ (Sperry, 1963). While the intervening 55 years have suggested a more complex mechanism including (but not limited to) such tags, no clear-cut cases of “Sperry” partner matching molecules that promote a direct, selective association of individual pre- and postsynaptic neurons had been identified. Due to a need to understand the molecular underpinnings of this process, though, a number of studies in recent years identified the Teneurins as key players in the partner matching step at the *Drosophila* neuromuscular junction (NMJ) and in the olfactory system (Mosca and Luo, 2014; Mosca et al., 2012). As such, a tempting conclusion is that in both the central and peripheral nervous systems, the Teneurins provide the strongest case to date for a Sperry molecule participating in these events.

In the relatively simple setting of the *Drosophila* NMJ, a single hemisegment contains 34 motoneurons that must each identify their appropriate muscle target among 31 options (Keshishian et al., 1993; Nose, 2012). What would a partner matching molecule look like in this situation? It would have to be expressed in both the presynaptic motoneuron and the postsynaptic muscle. Further, it would have to be expressed in a limited subset of motoneuron::muscle pairs – widespread expression would suggest it’s necessary for all connections, not specific ones. The first hypothesis that the Teneurins could serve this role at the NMJ came from expression studies. Though a basal level of Ten-m expression occurs in all larval muscles, two motoneuron::muscle pairs, those of muscle 3 and muscle 8, specifically express elevated levels of Ten-m (Mosca et al., 2012). To determine whether this expression was related to their function, work focused on altering Ten-m expression to better understand its function (Mosca et al. 2012). Knockdown of *ten-m* expression in muscle 3 and its innervating neuron (where it is normally highly expressed) increases the failure rate of innervation. The same failure occurs whether *ten-m* was knocked down in only the neuron or the muscle (Mosca et al. 2012), suggesting that both muscle 3 and its motoneuron require *ten-m* to properly match (i.e., both pre- and postsynaptic partners). This finding supports the idea of a homophilic interaction between pre- and postsynaptic Ten-m. This homophilic specificity was further elucidated by experiments showing that *ten-a* knockdown lacked such defects and Ten-a overexpression could not compensate for loss of Ten-m with respect to matching (Mosca et al., 2012).

Intriguingly, not only is Ten-m required for partner matching at the NMJ, it can also instruct matching between cells that do not normally connect. At muscles 6 and 7 in the developing larva, 40% of the boutons from the same motoneurons form on muscle 7 while the remainder form on muscle 6 (Johansen et al., 1989). However, the misexpression of Ten-m only in muscle 6 (but not 7) and both of their accompanying motoneurons, shifts the balance of connections to predominantly favor muscle 6 (Mosca et al., 2012). This suggests that expression of *ten-m* in cells that do not normally express it in high levels can direct partner-matching, similar to situations where *ten-m* expression is reduced. Interestingly, this was not recapitulated with Ten-a, suggesting not only a homophilic interaction between Ten-m that was instructive, but that there is some level of specificity regarding Ten-m over Ten-a. Therefore, the presynaptic level of Ten-m must be equivalent to the postsynaptic level of Ten-m for partner-matching to occur correctly.

This mechanism between pre- and postsynaptic targets suggested the first tenet of Teneurin function: transsynaptic interaction leading to partner matching, here, a homophilic interaction. But how do the Teneurins mediate this? What functions downstream of teneurin::teneurin interaction? A tantalizing possibility comes from work done to characterize the role of Teneurins in motor axon growth cone guidance (Zheng et al., 2011). Loss of *ten-m* caused aberrations in fasciculation that resulted in inter-segmental nerves moving to incorrect regions of the NMJ while ectopic overexpression of *ten-m* in the epidermis also caused axon migration defects. These defects were phenocopied by mutations in *cheerio*, the *Drosophila* homologue of the cytoskeletal protein filamin (Zheng et al., 2011). Filamin and Ten-m also interact physically (Zheng et al., 2011), suggesting that the Teneurins may control how neurons move and match with targets through interaction with the cytoskeleton. The defects in fasciculation caused by altering *ten-m* levels could be the same as the previously discussed defects with partner-matching after Teneurin perturbation. Furthermore, because Ten-m interacts with filamin, and mutations to filamin cause similar defects, partner-matching may very well be mediated by the reorganization of the cytoskeleton by a Ten-m/filamin complex. This would suggest a second tenet of Teneurin function: mediation of downstream function via interaction with the cytoskeleton. As such, additional research should explore this fascinating possibility that Teneurin-related partner matching requires modulation of the cytoskeleton.

Whether these mechanisms were selective for peripheral synapses or if they could also function in the central nervous system remained an open question. In the *Drosophila* olfactory system, however, there is a similar requirement for partner matching. Neurons in the *Drosophila* antennal lobe (Jefferis and Hummel, 2006), the first order processing center for olfactory information, must also match presynaptic axons of olfactory receptor neurons (ORNs) to postsynaptic dendrites of a cognate class of projection neurons (PNs). A genetic screen designed to identify potential partner matching molecules at this synapse (Hong et al., 2012) identified Ten-a and Ten-m as regulators of this process. In the olfactory system, all glomeruli have a basal level of both Ten-m and Ten-a, but some classes of neurons have elevated levels of either protein (Hong et al., 2012; Mosca and Luo, 2014). Specifically, certain matching ORN-PN pairs express elevated levels of the same Teneurin (Ten-a or Ten-m), reminiscent of Ten-m expression at the NMJ. Knocking down expression of both *teneurin* genes in both ORNs and PNs leads to mismatching between known partners, a phenotype that was also seen when both *ten-m* and *ten-a* are reduced in ORNs or PNs. More specific analysis revealed that the levels of Teneurin expression play a role in partner-matching. Knockdown of *ten-a* in PNs that normally have high Ten-a levels caused them to mismatch with ORNs that normally have low amounts of Ten-a. However, knockdown of *ten-a* in PNs that are naturally Ten-a low does not cause ORN mismatching, suggesting that PNs and ORNs must have similar levels of Ten-a to match correctly (Figure 1). A similar logic followed for ORN and PN pairs that expressed high levels of Ten-m (Hong et al., 2012). Altogether, these experiments suggested that partner-matching between PNs and ORNs occurs by a homophilic process in which both cells express either Ten-a or Ten-m at the same level. Similarly to the NMJ, this wiring could be mismatched by overexpressing *ten-m* or *ten-a* in a specific ORN or PN (Figure 1) that normally only has low levels of that teneurin, suggesting that these elevated levels of matching Teneurins can instruct partner matching (Hong et al., 2012). This again supports the notion of a conserved tenet where elevated Teneurin levels control partner matching between select cognate classes of ORNs and PNs.

**Figure 1.**
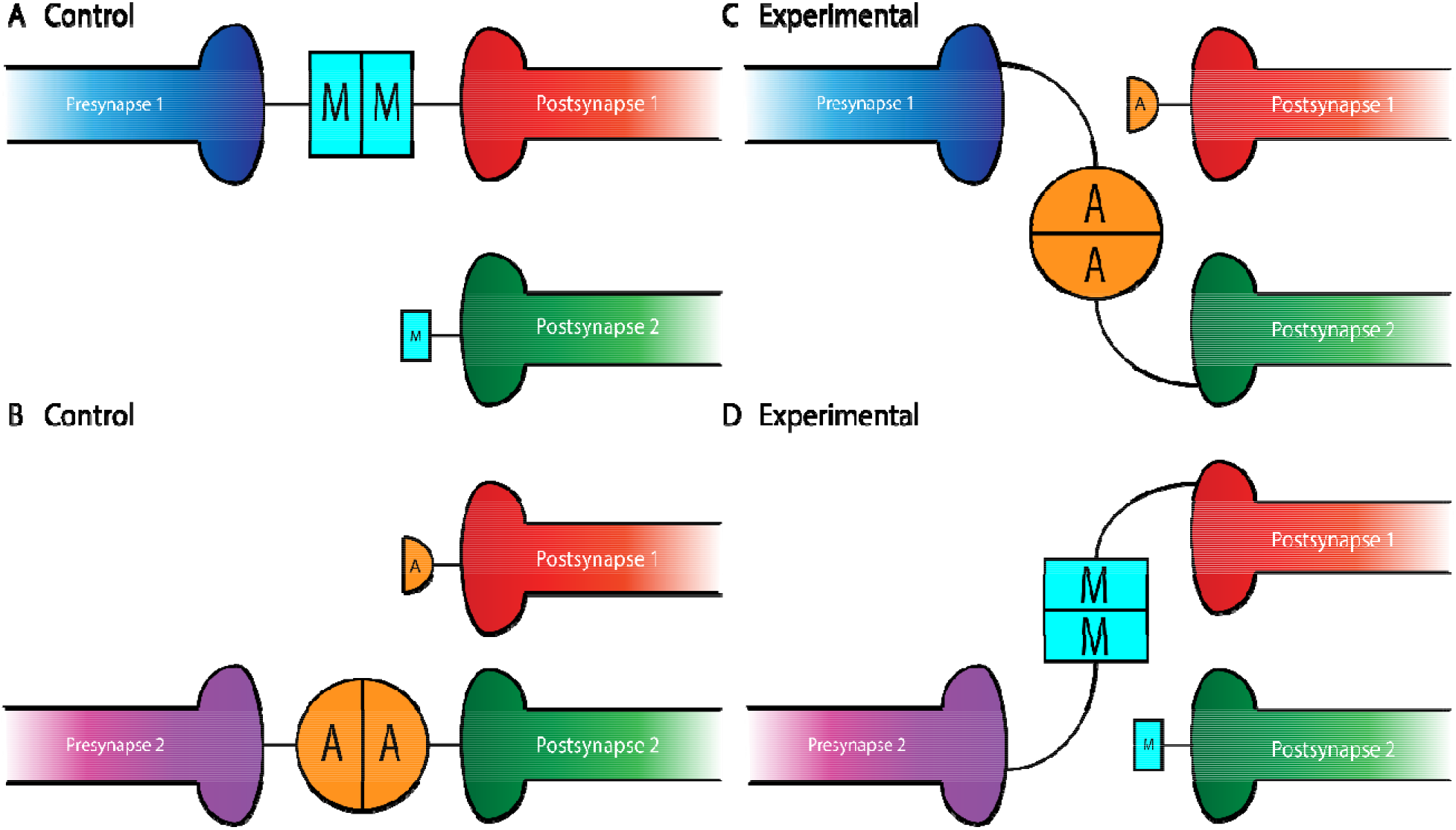
Teneurin-mediated partner matching in the CNS and PNS. Elevated levels of Teneurin expression control partner matching between distinct subsets of presynaptic neurons and their cognate postsynaptic partners. Normally, Presynapse 1 (either a motoneuron or ORN axon) with high levels of Ten-m matches with Postsynapse 1 (either a muscle or PN dendrite) expressing high levels of Ten-m, denoted here with a larger blue square (**A**). Presynapse 2 with high levels of Ten-a matches with Postsynapse 2 expressing high levels of Ten-a, shown above with a larger orange circle (**B**). In conditions where Teneurins are mis-expressed, alterations in wiring can occur. Experimental overexpression of Ten-a in the neuron of Presynapse 1 causes it to match with Postsynapse 2 which is also expressing high Ten-a instead of Postsynapse 1 (**C**). Overexpression of Ten-m in the neuron of Presynapse 2 causes it to match with Postsynapse 1 instead of its normal partner, Postsynapse 2 (**D**).

**Figure 2.**
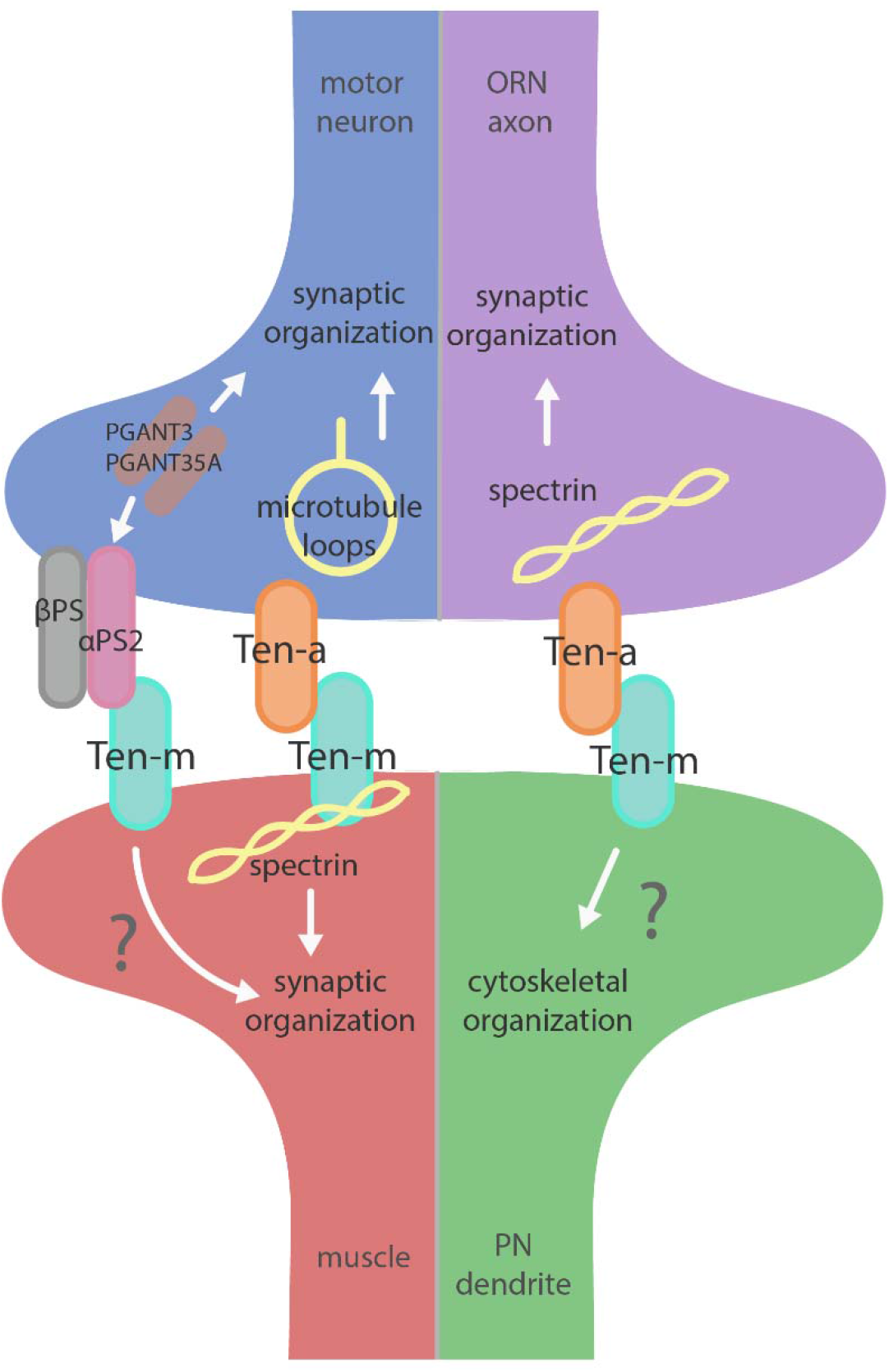
Teneurin-mediated synaptic organization in the CNS and PNS. Teneurins are involved in synaptogenesis in the PNS (left) and CNS (right). Both systems utilize transsynaptic heterophilic Teneurin interaction to instruct synaptic organization. At the NMJ, presynaptic Ten-a is involved in the formation of stable microtubule loops, promoting synaptic organization. Ten-m in the muscle interacts directly with spectrin, as well as potentially with αPS2 to mediate synaptic organization. In the CNS, however, presynaptic Ten-a functions with spectrin to promote synaptic organization. The role of postsynaptic Ten-m is unknown but may regulate downstream cytoskeletal components.

A number of open questions regarding the Teneurins and partner matching remain. Though some glomeruli follow a “Teneurin code” for expression and matching, others share overlapping expression and difference of phenotypic severity, suggesting partial redundancy between the Teneurins (Hong et al., 2012). The nature of this redundancy is not yet understood. In addition, little is known about other proteins involved in partner matching, as two Teneurins are not sufficient to pattern the entire antennal lobe. Recent work highlighted roles for Toll-6 and Toll-7 receptors (Ward et al., 2015), along with DIP/Dpr proteins (Barish et al., 2018), but a complete understanding remains elusive. The Teneurins may be part of a broader code involving a balance of additional proteins and their expression levels to determine the final correct partner match. Further still, it is unknown whether there is specificity for other cell types such as local interneurons or alternate connection modes such as dendro-dendritic connections between PNs, leaving an active area of study. Despite these unknowns, core tenets remain: Teneurins are required pre- and postsynaptically for partner matching and they do so in a homophilic fashion (Hong et al., 2012; Mosca et al., 2012) via elevated levels. This may occur via modulation of the cytoskeleton (Zheng et al., 2011) at both the NMJ and in the CNS, revealing a fascinating instance of mechanistic conservation. Their widespread expression in both invertebrate and vertebrate systems (Antinucci et al., 2013; Dharmaratne et al., 2012; Feng et al., 2002; Kenzelmann et al., 2008; Li et al., 2006; Mörck et al., 2010; Mosca, 2015; Zhang et al., 2018) and conservation of protein structure and mechanistic function (Jackson et al., 2018) suggests they may represent a general and versatile matching mechanism across synapse types and evolutionary taxa.

## SYNAPSE ORGANIZATION

Once a neuron has identified the correct synaptic partner, it must undergo synaptogenesis to form a functional and lasting connection. This is a complex process that involves multiple steps by which pre- and postsynaptic proteins align, synaptic machinery assembles, and the cytoskeletal components organize. Screens for synaptic molecules at the *Drosophila* NMJ have identified Teneurins as potential players in this process (Kurusu et al., 2008; Liebl et al., 2006) though their function in synaptic connectivity had not been elucidated until more recently. Work at the NMJ and in the olfactory system identified a conserved function for the Teneurins in synaptogenesis with a mechanism somewhat distinct from, though resembling, that of partner matching. In partner matching, the Teneurins use a homophilic transsynaptic interaction to partner match cells expressing the same, elevated levels of a particular Teneurin (Hong et al., 2012; Mosca et al., 2012). During synaptogenesis, however, Teneurins interact heterophilically and transsynaptically, with Ten-a being found mainly at the presynapse and Ten-m at the postsynapse (Mosca and Luo, 2014; Mosca et al., 2012). Because of this general role in synaptic organization, distinct from partner matching, all olfactory and muscular synapses show a basal level of Teneurin expression, while only select synapses participating in Teneurin-mediated partner matching show high levels of expression (Hong et al., 2012; Mosca et al., 2012). Still using the same family, the concept of Teneurin::Teneurin interaction is conserved, but the use of homo- versus heterophilic interactions allows for the diversification of processes. Further similar, the mechanism of Teneurin function regulating the cytoskeleton is also conserved in synaptic organization (Mosca and Luo, 2014; Mosca et al., 2012), an echo of the second tenet of Teneurin function suggested by partner matching and axon guidance studies. Here, we will explore the role of Teneurins in establishing synaptic connectivity in both the peripheral and central nervous systems and the evidence supporting these mechanisms.

The *Drosophila* NMJ is an ideal setting to study synaptic development in that it combines singular simplicity with powerful molecular genetics (Harris and Littleton, 2015). Studying the role of Teneurins at the NMJ opened a window into understanding their trans-synaptic role in synaptogenesis. There, Ten-a is expressed presynaptically, where it colocalizes with the active zone marker Bruchpilot and the periactive zone marker Fasciclin II (Mosca et al., 2012). Ten-m also shows low levels of presynaptic expression, but is predominantly expressed in the postsynaptic muscle, where it colocalizes with postsynaptic markers Dlg (the *Drosophila* homologue of PSD-95) and the cytoskeletal protein α-spectrin. Pre- or postsynaptic perturbation of Ten-a and Tenm (respectively) causes similar disruptions in synaptic structure and function, including a reduced number of synaptic boutons, defects in active zone apposition, general disorganization of synaptic components, impaired electrophysiological function, and defective vesicle cycling, many of which are reflected in severe locomotor impairment (Mosca et al., 2012). Taken together, these defects indicate a role for Teneurins in synaptic development. This suggested an extension of the first tenet of partner matching: a Teneurin::Teneurin interaction, but with a distinction that partner matching requires homophilic Teneurin interaction, and synaptic development requires heterophilic interaction of presynaptic Ten-a with postsynaptic Ten-m. Interestingly, tissue-specific removal of the presynaptic pool of Ten-m also results in a morphological phenotype, suggesting a presynaptic role (Mosca et al., 2012). Furthermore, postsynaptic Ten-m knockdown did not enhance the *ten-a* mutant phenotype, potentially suggesting presynaptic redundancy, or an additional postsynaptic receptor for presynaptic Ten-m. Overall, these experiments suggest a transsynaptic, heterophilic interaction between motoneuron-expressed, presynaptic Ten-a and muscle-expressed, postsynaptic Ten-m.

But what is the downstream mechanism for how the Teneurins mediate such effects? In addition to general defects in synaptic organization, the interruption of heterophilic Teneurin interaction at the NMJ also causes profound cytoskeletal disorganization. Teneurin perturbation causes a disruption of organized presynaptic microtubule loops and an increase in unbundled Futsch / MAP-1b staining, suggesting a deranged cytoskeleton (Mosca et al., 2012). Additionally, the loss of Teneurin signaling also causes a near complete loss of the postsynaptic spectrin cytoskeleton. As direct cytoskeletal disruption can serve as a common cause for many of the phenotypes observed following Teneurin perturbation, this lead to the hypothesis that Teneurins organize synapses via a link with the cytoskeleton. Indeed, Ten-m colocalizes with and physically interacts with α-spectrin in a complex (Mosca et al., 2012). As spectrin is a molecular scaffold which interacts with actin to form a network along the inside of the plasma membrane, this suggested that Ten-m may represent the link between the synaptic cytoskeleton and the membrane, further strengthening the hypothesis of direct cytoskeletal interaction with the Teneurins. Additionally, loss of postsynaptic spectrin does induce similar synaptic growth defects (Pielage et al., 2006), which is consistent with this hypothesis. Thus, Teneurins are involved in organizing synapses by way of ordering the cytoskeleton, as mediated through a Ten-m link between the synaptic membrane and α-spectrin (Mosca et al., 2012). This further supports the second tenet of Teneurin function: mediating their role in neuronal development via cytoskeletal modulation.

Though playing a critical role in synaptic organization, the Teneurins also cooperate with other cell surface proteins to construct a connection. Neurexin and the Neuroligins are transmembrane proteins that instruct synaptic development; phenotypes associated with their disruption include changes in bouton number and disorganization of active zones (Banovic et al., 2010; Li et al., 2007; Owald et al., 2012; Sun et al., 2011; Xing et al., 2014; Zhang et al., 2017). These results have considerable phenotypic overlap with perturbation of *ten-a* and *ten-m*, suggesting potential genetic or pathway interaction. The two instead operate in distinct but partially overlapping pathways: Neurexin / Neuroligin 1 largely control active zone apposition with minor effects on the cytoskeleton while the Teneurins largely control cytoskeletal organization and cooperate with Neurexin / Neuroligin1 to regulate active zone apposition. This reveals that there is a complex cooperation between cell surface proteins and a division of labor to ensure that synaptic contacts are properly organized.

Teneurins also show remarkable similarities in how they function in the central nervous system, as evidenced by examination of transsynaptic Teneurin signaling in the *Drosophila* olfactory system (Mosca and Luo, 2014). The olfactory system is valuable for studying synaptic development due to its well defined synaptic connections in the context of a complex circuit (Jefferis and Hummel, 2006). At ORN synapses, perturbations in Teneurin levels (either presynaptic Ten-a in the ORNs or postsynaptic Ten-m in the projection neurons) also impaired synaptic organization. The number of both presynaptic active zones and postsynaptic acetylcholine receptors are decreased when presynaptic *ten-a* or postsynaptic *ten-m* are knocked down. This is a strikingly similar phenotypic result as to the NMJ, suggesting conservation between the CNS and the PNS. The mechanism is also strikingly similar, as Ten-a and Ten-m interact heterophilically, and are found primarily at the pre- and postsynapse, respectively. Furthermore, the link with spectrin is also conserved between the two systems: Ten-a and spectrin function in the same genetic pathway to control central synapse number, the spectrin cytoskeleton in the antennal lobe is drastically reduced following Teneurin perturbation, and presynaptic spectrin is also important for achieving normal synapse number in ORNs. As in the peripheral nervous system, Teneurins function with the cytoskeleton to allow proper cytoskeletal organization for the formation of a robust synaptic architecture (Mosca and Luo, 2014). Thus, the second tenet of Teneurin function, downstream regulation of the cytoskeleton, is further conserved. Finally, a third conserved aspect links Teneurins with synaptic function. At the NMJ, *ten-a* mutants show reduced evoked postsynaptic potentials, impaired vesicle cycling, and reduced larval locomotion (Mosca et al., 2012). Restoring Ten-a expression to motoneurons partially rescues the locomotor phenotype, suggesting a presynaptic function for Ten-a in regulating function. In the CNS, olfactory function can be measured by behavioral response: basic function can be assayed by the performance of flies in a modified olfactory trap (Larsson et al., 2004; Min et al., 2013; Mosca et al., 2017; Potter et al., 2010) using apple cider vinegar (ACV) as an attractive odorant source. Control flies are nearly uniformly attracted to ACV (Figure 3) – impaired attraction can be indicative of synaptic defects, as seen when the synaptic organizer LRP4 is removed specifically from ORNs (Mosca et al., 2017). *ten-a* mutants have significantly impaired ACV attraction (Figure 3), suggesting that *ten-a* is required for normal olfactory function (as it is required for normal neuromuscular function). Studies in the whole-animal mutant, however, do not determine where Ten-a functions to regulate function. To address this, we restored *ten-a* expression only to adult ORNs using *pebbled-GAL4* and found that this partially rescued the loss of olfactory attraction (Figure 3). This suggests that presynaptic Ten-a mediates normal function at olfactory synapses, again similar to the NMJ. Thus, the conservation of a role for Teneurins in promoting normal presynaptic function represents a third tenet of Teneurin mechanisms that span the olfactory and neuromuscular systems. There are, however, some variations on the organizational theme between the CNS and the PNS. Interestingly, the spectrin interaction seems to occur presynaptically in the CNS but postsynaptically at the NMJ. Also, a mild phenotype is present when Ten-m is knocked down at the presynaptic NMJ, indicating a minor presynaptic role, but no such phenotype is observed in the CNS (Mosca et al., 2012, Mosca and Luo 2014). This indicates perhaps that though the broad mechanisms may be conserved, certain elements differ, perhaps owing to the differing complexity and biological role for each synapse. This offers an interesting way to diversify a conserved mechanism – with mild adjustments to allow for different kinds of synapses. Teneurins may also promote postsynaptic cytoskeletal organization in the CNS, but that interaction has yet to be identified.

Beyond the spectrin cytoskeleton, additional work has suggested broader conservation. Teneurins also regulate the cytoskeletal proteins adducin and Wsp (Mosca et al., 2012) and can further interact with integrins via αPS2, a synaptic integrin receptor (Graner et al., 1998). At the *Drosophila* NMJ, knockout of alpha-N-acetylgalactosaminyltransferases (PGANTs), proteins which regulate integrins, led to decreased levels of αPS2 as well as Ten-m (Dani et al., 2014). Ten-m at the presynapse may be involved in cell adhesion through interaction with αPS2, causing the mild phenotype observed when Ten-m is knocked down only in neurons. Future work on the roles of Teneurins will further determine their effectors and how these factors serve to instruct synaptic connectivity via regulation of the cytoskeleton.

Much like partner matching, the *Drosophila* Teneurins play a critical role in synaptic organization. Further, their function is conserved in the CNS and PNS and also, in a mechanistic fashion by 1) a transsynaptic interaction and 2) a regulation of the downstream cytoskeleton. However, certain distinctions make the organizational process unique from partner matching. Here, basal levels of Teneurins mediate synaptic organization through a heterophilic transsynaptic interaction: Ten-a is predominantly presynaptic while Ten-m is postsynaptic. Further, the Teneurins are relatively unique among synaptic organizers in that their main role is to mediate cytoskeletal components. The remarkable evolutionary conservation present within these systems indicates the importance of Teneurins in their various roles. Further work is needed to examine the specific functions of Teneurins in regulating synaptic connectivity, but the widely conserved mechanisms already observed in Teneurin function promise the advantage of continued study across systems and synapses (Mosca, 2015).

## FUTURE DIRECTIONS

The complex series of events that underlie neuronal development have distinct molecular, temporal, and spatial requirements that safeguard their fidelity. To ensure evolutionary economy, these events can be coordinated through reuse of molecular cues and mechanisms. These mechanisms are conserved from peripheral to central synapses in *Drosophila*; work has also shown similar roles in mammalian nervous systems for wiring and synapse organization (Berns et al., 2018; Dharmaratne et al., 2012; Leamey et al., 2007; Mosca, 2015), suggesting mechanistic conservation across multiple species as well. As such, the Teneurins are an evolutionary constant, working at multiple levels to ensure nervous system development. In our current understanding, however, there is much left to learn about how Teneurins regulate nervous system development. Recent work especially has highlighted the interplay between Teneurins and Latrophilin in mammalian synapse organization (Boucard et al., 2014; Li et al., 2018; Silva et al., 2011; Vysokov et al., 2016) and in behavioral regulation through TCAP, the Teneurin C-terminal Associated Peptide (Woelfle et al., 2015, 2016). In *Drosophila*, potential interactions between the Teneurins and Latrophilin have not been studied. The *Drosophila* genome possesses a single Latrophilin homologue, *dCirl* (Scholz et al., 2015). *dCirl* is expressed in larval chordotonal neurons and is required for mechanosensation and larval locomotion (Scholz et al., 2015). In these neurons, *dCirl* functions to reduce cAMP levels in response to mechanical stimulation (Scholz et al., 2017). Whether these functions involve Teneurins remains an open question. There is likely not complete overlap between Teneurins and dCIRL, as *dCirl* mutants do not phenocopy the synaptic defects associated with *ten-a* / *ten-m* perturbation (T. Mosca, unpublished observations). This does not, however, address potential redundancy in the genome with other GPCRs or orphan receptors, so more directed study is needed. As Teneurins are also thought to interact with other cell surface receptors (Mosca et al., 2012) and adhesion molecules (Dani et al., 2014), it is increasingly likely that Teneurins represent a nexus for receptor interaction, suggesting that a number of players are yet to be discovered.

One key unanswered question involves the role of presynaptic Ten-m at the NMJ. Though predominantly postsynaptic at the NMJ (Figure 4A), Ten-m also localizes presynaptically in motoneurons (Mosca et al., 2012); this contribution is revealed when Ten-m is removed specifically from the muscle using RNAi (Figure 4B). Presynaptic knockdown of Ten-m results in a modest reduction in synaptic bouton number (Mosca et al., 2012). However, as Teneurins and Integrins all promote synaptic maturation (Lee et al., 2017; Mosca et al., 2012), and Ten-m may interact with integrins (Dani et al., 2014), this raises the possibility that *ten-m* may contribute to activity-dependent synaptic remodeling (Ataman et al., 2008; Piccioli and Littleton, 2014; Xiao et al., 2017). At the *Drosophila* NMJ, acute spaced stimulation using high K^+^ induces activity-dependent sprouting in the form of “synaptopods” (Ataman et al., 2008). These synaptopods form in as little as 15-20 minutes and contain Ten-m (Figure 4C). This suggests that Ten-m is one of the first components of these nascent neurite branches. Ten-m is present even before synaptic vesicles appear, which are among the earliest components visible in ghost boutons (Ataman et al., 2008). This raises the possibility that Ten-m could promote synaptic maturation and activity-dependent growth. Further experiments will be needed to directly test this hypothesis but could more deeply connect presynaptic Tenm, neuromuscular growth, and integrins.

Further, our knowledge of downstream Teneurin effectors remains incomplete. At the *Drosophila* NMJ, neither reduced Neurexin / Neuroligin signaling (Banovic et al., 2010; Li et al., 2007) nor a loss of spectrin (Pielage et al., 2005, 2006) can account for the entire cadre of synaptic phenotypes associated with *teneurin* perturbation (Mosca et al., 2012). This suggests that additional downstream mechanisms exist to mediate Teneurin function. This could be through additional cytoskeletal proteins, as in *C. elegans* (Mörck et al., 2010). A more thorough understanding of how Teneurins engage partner matching, either by downstream mechanisms or interaction with other cell surface proteins is also poorly understood. Whether Teneurins interact with axon guidance molecules and cell surface receptors, as in *C. elegans* (Mörck et al., 2010) is a distinct possibility. Approaches to understand Teneurin-interacting proteins will be essential to understand the different ways they regulate their diverse functions.

Finally, a core question intrinsic to the Teneurins remains. As we understand, Teneurins use multiple interactive mechanisms to enable development: homophilic interactions match and maintain partners while heterophilic interactions organize synaptic connections. This must mean that, at the same connection, both homophilic and heterophilic interactions exist simultaneously. As these distinct pairs have distinct goals, how does a cell interpret which interaction is happening for a particular Teneurin molecule? For example, ORNs that use elevated Ten-a to match their cognate PNs also use basal levels of presynaptic Ten-a to organize their output synapses by interacting with postsynaptic Ten-m. Therefore, these ORNs simultaneously have homophilic and heterophilic interacting Ten-a molecules. How are these distinguished? Are certain downstream interactors only expressed at certain developmental times? This way, the downstream effectors specific to partner matching would only appear during times of neuronal wiring and be downregulated by the time synaptic formation, organization, and maintenance take over as the predominant processes. As partner matching and synapse formation can be separated by as much as 24-48 hours in the developing olfactory system (Jefferis and Hummel, 2006) or by as much as 4-6 hours at the developing NMJ (Broadie and Bate, 1993), this is a reasonable hypothesis. However, if this is not the case, it could be that the mechanism is more intrinsic to the Teneurin protein. If there was a fundamental difference between a Ten-a::Ten-a and a Tena::Ten-m interaction, this could result in conformational changes that only allowed binding of specific downstream molecules. One hypothesis is that this fundamental difference could come from tension (Mosca, 2015). The NHL domain present in the extracellular domain of Teneurins is thought to mediate interaction in *trans* (Beckmann et al., 2013). Homophilic NHL domain interactions display stronger adhesive forces than heterophilic (Beckmann et al., 2013): if this tension can be “read out” by the cell, it could recruit different downstream effector molecules depending on the transsynaptic partner of that Teneurin. This could enable a mechanism to distinguish homophilic from heterophilic Teneurin interactions when both may exist in the same small synaptic region. With more recent structural information about the Teneurins (Li et al., 2018), more directed hypotheses about interaction can now be explored. Beyond an intrinsic tension mechanism, more recent work showed that splice variants of Ten-3 in mouse can regulate cell-to-cell adhesion, potentially affecting neuronal wiring (Berns et al., 2018). Thus, there are multiple options for intrinsic ways that Teneurins could distinguish themselves depending on partners and interactions. Future work will be needed to dissect both the intrinsic and extrinsic mechanisms that enable Teneurins to function so broadly. Work over the last decade has cemented the Teneurins as essential regulators of neuronal development, functioning via related mechanisms in steps ranging from the initial elements of neuronal development in axon guidance, seeing the developmental process through to the end with functions in synaptic organization. Science will take the next bold steps forward from that foundation, venturing out to determine how these core cell surface proteins mediate downstream function, and moving closer to understanding the intricacies and complexities of neuronal development.

## ACKNOWLEDGMENTS

We would like to thank members of the Mosca Lab for stimulating discussions and camaraderie. This work was supported by awards to TM from the National Institutes of Health (R00-DC013059), the Commonwealth of Pennsylvania CURE Fund (SAP 4100077067), and the Alfred P. Sloan foundation.

